# PG-SCUnK: measuring pangenome graph representativeness using single-copy and universal K-mers

**DOI:** 10.1101/2025.04.03.646777

**Authors:** Cumer Tristan, Milia Sotiria, Alexander S. Leonard, Pausch Hubert

**Author notes:** Contact: Tristan Cumer –.

## Abstract

**Background:** Pangenome graphs integrate multiple assemblies to represent non-redundant genetic diversity. However, current evaluations of pangenome graphs rely primarily on technical parameters (e.g., total length, number of nodes/edges, growth curves), which fail to assess how effectively the graph represents homologous stretches across the integrated assemblies.

**Results:** We introduce a novel method to quantitatively assess how well a pangenome graph represents its integrated assemblies. Our method quantifies how many single-copy and universal k-mers from the source assemblies are uniquely and completely represented within the graph nodes. Implemented in the open-source tool PG-SCUnK, this approach identifies the fractions of unique, duplicated, and split k-mers, which correlate with short read mapping rates to the pangenome graph.

**Conclusions:** Insights provided by PG-SCUnK facilitate the selection of appropriate parameters to build optimal pangenome graphs.

**Availability and implementation:** A bash implementation of the PG-SCUnK workflow is freely available under the GNU GPLv3 license at https://github.com/cumtr/PG-SCUnK/.

## 1. background

Pangenome graphs have emerged as a promising data structure to overcome the limitations of linear reference genomes, which fail to capture genetic diversity thereby introducing mapping biases particularly for diverged genomes (Günther and Nettelblad 2019; Martiniano *et al*. 2020; Lin *et al*. 2024). Pangenome graphs integrate multiple assemblies and represent their diversity in a single graph (Abel *et al*. 2020). In such a graph, each node corresponds to a DNA sequence, with input assemblies represented as paths and edges connecting nodes along these paths. Conserved sequences appear as nodes shared across all paths, while variant sites form bubbles or snarls composed of different nodes and edges (Paten *et al*. 2018). Pangenome graph references can reduce read mapping and variant genotyping biases (Garrison *et al*. 2018; Sirén *et al*. 2021). Consequently, these graphs enable accurate genotyping across a wide range of variant types, from single nucleotide polymorphisms to large structural variants (SVs) (Hickey *et al*. 2020).

Various statistics are available to assess the quality, contiguity and completeness of linear assemblies (Li and Durbin 2024). Assembly contiguity is often evaluated with technical metrics like total length or contig L50 and N50, while assembly completeness is commonly assessed through the presence (complete or fragmented), absence, or duplication of highly conserved genes (i.e. using BUSCO scores (Simão *et al*. 2015)). For pangenome graphs, existing metrics primarily describe the graph’s structure without addressing its biological relevance. Widely used summary statistics report the total nucleotide length, the number of nodes, edges or paths (Guarracino *et al*. 2022), as well as node coverage and pangenome growth (Parmigiani *et al*. 2024). Complex genomic regions (such as centromeres) are often poorly represented in pangenome assemblies and can disproportionately inflate these metrics, thereby complicating their biological interpretation (Milia *et al*. 2024). Recent work evaluated graph quality by re-aligning either the original long read sequences (Liao *et al*. 2023) or source assemblies (Leonard *et al*. 2023) to the pangenome graph. However, none of these approaches comprehensively evaluate how accurately and efficiently the pangenome graph represents the underlying assemblies.

Here, we use single-copy and universal sequences to measure how well a graph represents homologous sequences of the source assemblies. We assume that k-mers found exactly once in an assembly (referred to as single copy) and present across all source assemblies of a pangenome (thus universal) are orthologous, thereby should be integrated uniquely in the nodes of the graph. Single-copy and universal k-mers (SCUnKs) from the source assemblies are classified as unique if they appear only once in full length, duplicated if they occur multiple times, and split if they are fragmented across different nodes in the evaluated pangenome graph. We have developed a tool to calculate these three metrics and demonstrated their utility in constructing pangenome graphs that achieve high mapping rates.

### 2. Implementation

We developed PG-SCUnK, a workflow designed to assess pangenome graph quality using SCUnKs derived from the input assemblies. PG-SCUnK categorizes these k-mers into three types depending on their frequency in the pangenome graph nodes:

- Unique: K-mer present only once and in full within the nodes.
- Duplicated: K-mer present multiple dmes in full within the nodes.
- Split: K-mer absent from the nodes, thus fragmented across nodes.

The PG-SCUnK workflow is as follows (Fig. 1):

**Figure 1.**
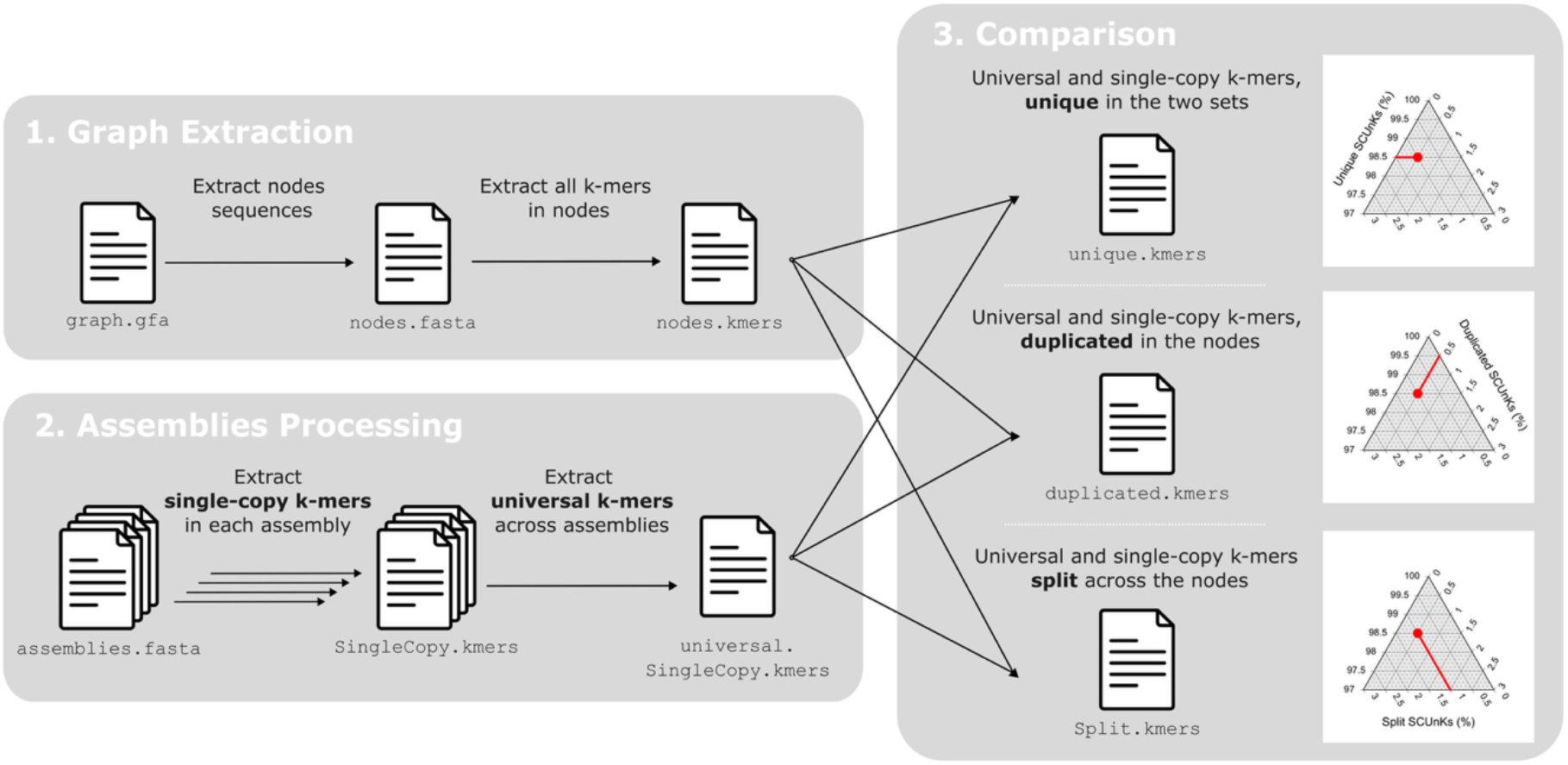
PG-SCUnK workflow. Grey shaded boxes represent the three main steps of the workflow. Captions on the right depict how PG-SCUnK results for a pangenome graph can be represented on a ternary plot, where each point represents the three metrics calculated by PG-SCUnk that sum to 100%. This example graph has 98.5% of the SCUnKs identified as unique, 0.5% identified as duplicated and 1% as split.

### Graph extraction

Sequences are extracted from all nodes in the pangenome graph and every k-mer is identified and its frequency of occurrence counted.

### Assemblies processing

K-mers present only once within each assembly used to construct the graph (single-copy k-mers) are identified. Single-copy k-mers that are present in all the input assemblies are referred to as SCUnKs.

### Comparison

The set of SCUnKs from the input assemblies is compared with those integrated into the graph to identify unique, duplicated, or split k-mers.

The PG-SCUnK workflow is implemented as a Bash script that automates these three steps, leveraging the KMC software (Kokot, Długosz and Deorowicz 2017) for efficient k-mer extraction and comparison.

## 3. Results and discussion

We evaluated the performance and output produced by PG-SCUnK on pangenome graphs built for different species using two widely used construction methods. Our evaluation included a cattle pangenome graph generated for this study (initially published by Milia *et al*. 2024; see supplementary material for details) and four previously published pangenome graphs: two human pangenome graphs (Liao *et al*. 2023) that were constructed using either *PGGB* (Garrison *et al*. 2023) or *Minigraph-Cactus* (Hickey *et al*. 2024), a pangenome graph for finches (Fang and Edwards 2024) and one for grapevine (Liu *et al*. 2024). See Supplemental Table S1 for detailed information about the graphs.

### Runtime and performances

The analysis of k-mer profiles of pangenome graphs with PG-SCUnK is both time- and memory-efficient (Table 1). For example, processing a graph containing 90 haplotype assemblies of a single human chromosome on a single thread required between 19 and 138 minutes per chromosome. Runtime scaled linearly with chromosome size (Fig. S1). Maximum memory usage ranged from 3.6 to 22.1 GB and significantly correlated with graph complexity (Fig. S1).

**Table 1.** Computational performance of PG-SCUnK. For each pangenome graph considered, the number in brackets reports the number of haplotypes included in the graph. For CPU time and peak memory usage, the number represents the median value across the different chromosomes while the number in brackets reports the minimum and maximum. All results presented here are for a k-mer size of 100.

| ORAGNISM | SOFTWARE | CPU TIME (S) | PEAK MEM. USAGE (GB) | REFERENCE |
| --- | --- | --- | --- | --- |
| CATTLE (21) | PGGB | 609 (314, 1614) | 7.42 (3.53, 14.59) | This study |
| HUMAN (90) | PGGB | 2946.5 (1156, 8251) | 11.94 (3.66, 17.85) | Liao *et al.* 2023 |
| HUMAN (90) | Minigraph-Cactus | 3868.5 (1501, 7102) | 14.88 (9.32, 22.07) | Liao *et al.* 2023 |
| FINCH (37) | PGGB | 277 (177, 1277) | 1.24 (0.69, 10.7) | Fang and Edwards 2024 |
| GRAPEVINE (29) | PGGB | 289 (258, 374) | 2.4 (1.61, 3.43) | Liu *et al.* 2024 |

### PG-SCUnK across diverse taxa

PG-SCUnK categorizes the SCUnKs into unique, duplicated, and split, thereby revealing apparent differences among the evaluated pangenome graphs (Fig. 2 & S2). We observed a high proportion of unique SCUnKs ranging from 97.93 to 99.04 in the human pangenome graph built with *PGGB*, and from 99.11 to 99.42% in the cattle graph. Only a small fraction of the SCUnKs were either duplicated (duplication rates of 0.03-0.94% and 0.06-0.34% for humans and cattle respectively) or split (split rates of 0.89-1.47% and 0.49-0.65%). Human chromosomes displayed some heterogeneity in the PG-SCUnK scores, and this pattern was consistent across both construction methods (i.e. *PGGB* and *Minigraph-Cactus*, Fig. S3). Much greater PG-SCUnK scores variability was observed for the finch and grapevine graphs; the finch chromosomal graphs showed a wide range in unique and duplication rates (43.6 to 95.08% for unique and 2.32 to 55.90% for duplicated) coupled with low split rates (0.41 to 2.88%). The grapevine graphs exhibited variable unique and split rates (79.92 to 96.39% and 3.42 to 20.08% respectively) with low duplication levels (0 to 0.52%). Further research is needed to explore if technical factors (e.g., differences in graph construction parameters) or biological factors (e.g., assembly characteristics such as GC content, repeat regions or completeness) contribute to the observed heterogeneity between and within pangenomes. While exploring the reason behind the variability of each specific case is beyond the scope of this study, we provide a companion script (FindSCUnKsRegions.bash) to map unique, duplicated, and split SCUnKs back to the genome. This allows users to identify the genomic regions where these SCUnKs occur which may provide useful to investigate why certain SCUnKs are duplicated or split (Figure S4-S7). Nevertheless, our results demonstrate that PG-SCUnK efficiently exposes how well pangenome graphs represent homologous sequences from the assemblies across the tree of life.

**Figure 2.**
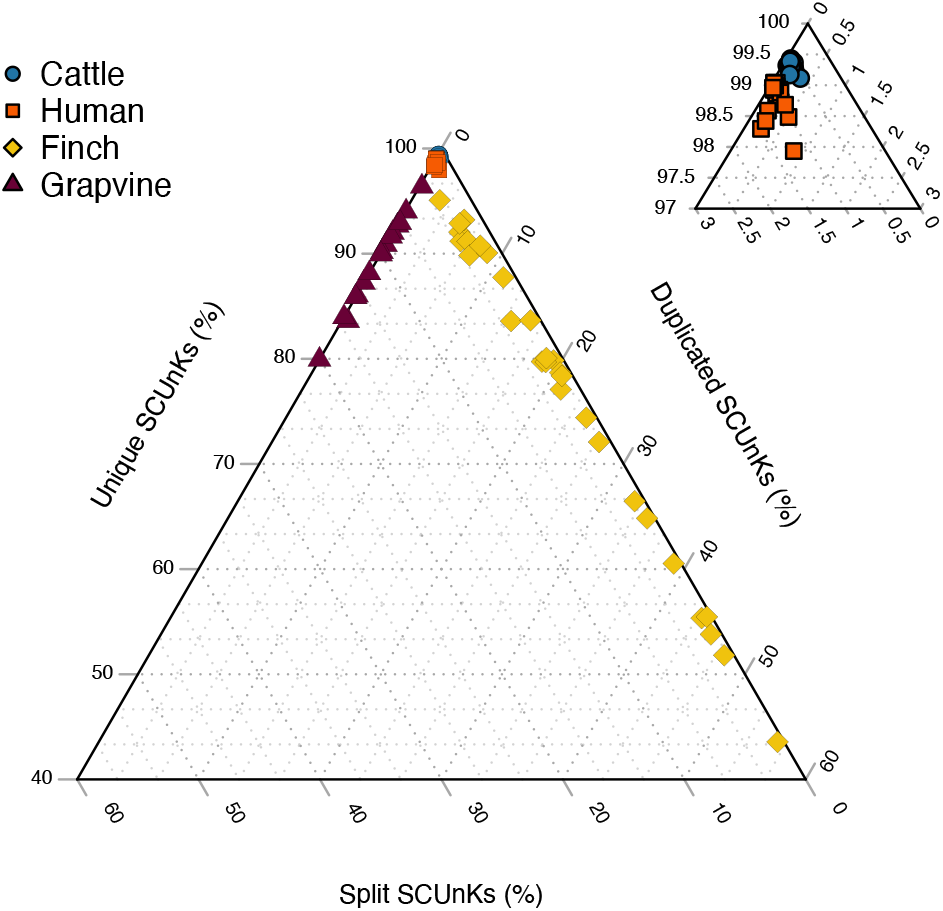
PG-SCUnK scores for various pangenome graphs. The ternary plot presents the PG-SCUnK scores for publicly available pangenome graphs of four species (cattle, human, finch and grapevine). The right panel depicts a detailed view of the top corner of the main ternary plot.

### Effects of k-mer size

We investigated how k-mer size (31, 51, 71, 100 [default], 111, and 211) affects the portion of the genome considered by PG-SCUnK and the inferred scores. Our results revealed striking differences in the absolute proportion of k-mers identified as SCUnKs, ranging for example from 69.3% (-k 31) to 19.7% (-k 211) in humans and from 8.8% % (-k 31) to 0.1% (-k 211) in finches (Fig. S8). This variation may be driven by species-specific heterozygosity, as greater genetic diversity reduces the likelihood of a k-mer being universal (genome wide heterozygosity between 0.0013 to 0.0016 in humans (Auton *et al*. 2015) and 0.003 to 0.005 in finches (Shultz *et al*. 2016)). Across all tested graphs, the proportion of SCUnKs decreased with increasing k (Fig. S8), likely because larger k-mers are more prone to disruptions by polymorphisms. Despite these variations, k-mer size has little to no effect on PG-SCUnK scores for k ≤ 111 (Fig. S9 - S12): the correlation between PG-SCUnK scores ranged from 0.935 to 0.999 in cattle, 0.992 to 1 in humans, 0.974 to 1 in finches and 0.967 to 0.998 in grapevine. These findings indicate that while k-mer size influences the fraction of the genome considered, its impact on PG-SCUnK scores remains minimal.

### PG-SCUnK score varies depending on the construction parameters

We explored the usefulness of PG-SCUnK for optimizing graph construction assuming that SCUnKs should be found in their full length and unique in an optimally built graph (i.e. maximizing the rate of unique SCUnK). We generated multiple graphs with *PGGB* for bovine chromosome 13 using varying parameter sets and calculated PG-SCUnK scores for the resulting graphs (Fig. 3A). Adjustments to the segment-length parameter (-s) produced distinct effects: low values increased the split rate (indicative of an overly compact graph), intermediate values maximized uniqueness (with a maximum value obtained for -s 10k), and higher values led to increased duplication rates. A similar pattern was observed for the percent-identity parameter (-p), with an optimization of the unique rate at intermediate values. The minimum match length parameter (-k) has no impact on the PG-SCUnK scores at low values but results in a sharp increase in duplication and split rates at higher values. These results demonstrate that PG-SCUnK can be used to guide parameter selection during pangenome graph construction and can aid building high quality reference graphs.

**Figure 3.**
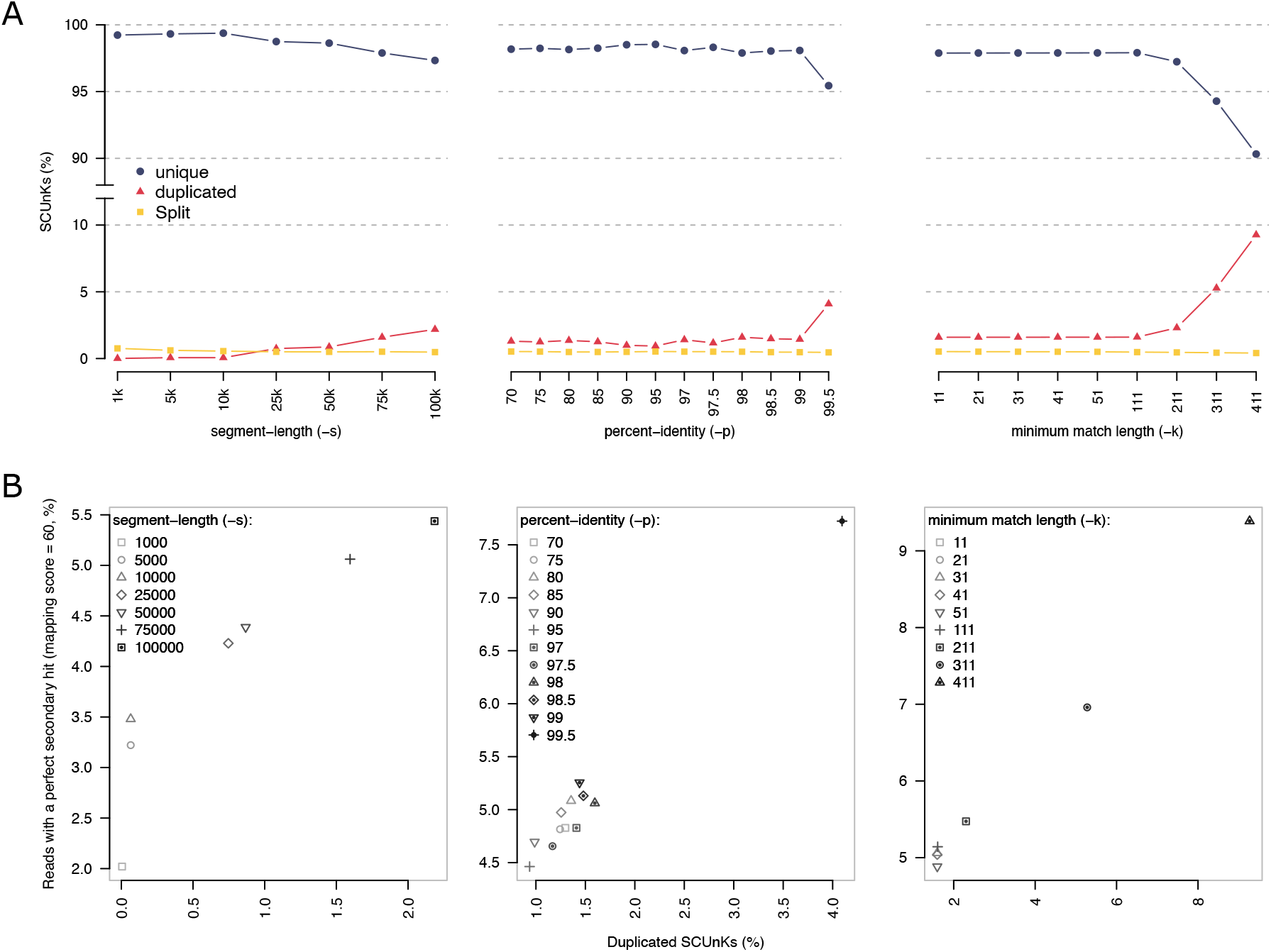
PG-SCUnK scores varies depending on parameter choice and predict the mappability of the graph. (A) PG-SCUnK scores across a wide range of parameters used during the building of a cattle pangenome graph for chromosome 13. For each varying parameter, the two others were fixed to the values used for the entire genome (fixed parameters: -s 75k, -p 98, k 31). (B) Relation between the duplication rate of the SCUnKs identified by PG-SCUnK and the proportion of reads with secondary alignments with a mapping quality of 60 across the range of the graph presented in (A).

### PG-SCUnK scores predict the mappability of a graph

We evaluated the ability of PG-SCUnK to predict the mappability of a pangenome graph, based on the hypothesis that elevated levels of duplicated or split SCUnKs would compromise read mapping accuracy.

Using the bovine pangenome graph constructed with various parameters, we simulated short reads from the source assemblies, mapped them to the graph, and assessed two key metrics: (i) the proportion of reads mapping with a perfect score (mapping quality of 60), and (ii) the proportion of those reads with a secondary mapping of equivalent quality.

A high proportion of perfectly mapped reads supports the graph’s utility as an unbiased reference. As expected, across all parameter settings, the proportion of simulated reads mapping with a quality score of 60 remained consistently high. For the segment-length parameter (-s), the mean proportion ranged from 92.4% to 93%, except at a segment length of 1 kbp, where it dropped to 83.7% (Fig. S13). For the percent-identity parameter (-p), values ranged from 92.2% to 93.6%, with a notable increase to 98.2% at 99.9% identity (Fig. S13). For the minimum match length parameter (-k), the mapping rate varied between 92.6% and 96.2%, with the lowest rate observed at k = 51 (Fig. S13). Additionally, we observed a negative relationship between the proportion of split SCUnKs and mapping rate: graphs with the highest mapping rates had the lowest split SCUnK proportions (Fig. S14).

To assess mapping confidence, we examined the proportion of reads with a secondary alignment of equivalent quality (i.e., both primary and secondary mapping quality = 60). A low rate indicates confident, unique mapping, while a high rate suggests ambiguity. This metric revealed substantial differences across graph-building parameters. The proportion of duplicated mappings increased from 1.4% to 4.6% with varying segment-length values (-s; Fig. 3B, S13), ranged from 3.9% to 6.7% for percent-identity (-p), and spiked to 80.5% at p = 99.9 (Fig. 3B, S13). For minimum match length (-k), the proportion ranged from 4.1% to 8.5%, with a steep rise at higher values (Fig. 3B, S13). These trends mirrored the duplication rates captured by PG-SCUnK scores, confirming a clear relationship between duplicated SCUnKs and secondary mapping rates (Fig. 3B).

Consistent with our single-chromosome observations, a high proportion of reads mapped perfectly across all chromosomes in each organism: 98.6% for humans, 99.1% for cattle, 99.2% for finches, and 97.4% for grapevine (Fig. S15). This proportion was negatively correlated with the fraction of split SCUnKs across organisms (Fig. S16), though intra-organism chromosomal comparisons showed more variability (Fig. S17). The rate of secondary alignments of equivalent quality revealed marked differences between species, with mean values of 0.023 for humans, 0.025 for cattle, 0.15 for finches, and 0.027 for grapevine (Fig. S18), alongside some heterogeneity within each species’ chromosomal graphs. Across all organisms, the fraction of duplicated SCUnKs was positively correlated with the rate of perfect secondary alignments (Fig. S19), a trend largely driven by the finch pangenome but consistently observed across species (Fig. S20).

Taken together, these results demonstrate that PG-SCUnK scores predict graph mappability and offer a practical framework for selecting optimal reference graphs for accurate short-read mapping.

### PG-SCUnK scores outperform existing metrics

To the best of our knowledge, there are currently no methods or metrics that formally assess how well a graph reflects its constituent assemblies. However, some existing technical metrics can still be useful to assess the properties of pangenome graphs. One of these metrics is total inflation, which is defined as the ratio of the graph’s total nucleotide length to that of a single assembly (typically the highest-quality or reference assembly).

We evaluated how the total inflation score can predict graph mappability and compared it with the PG-SCUnK score (Figure S21). Our results show that, within each organism, inflation correlates with the relative mappability of the chromosomal graph; specifically, higher inflation scores are associated with increased duplication rates. However, absolute inflation values varied substantially across organisms. For instance, certain cattle graphs exhibited inflation scores twice as high as the maximum observed in finches, despite having less than a third of the duplication rate. In contrast, the PG-SCUnK score provides a more absolute measure of mappability by relating the SCUnK duplication rate to the probability of a read having a secondary hit with a mapping quality of 60.

Overall, these findings suggest that while total inflation may serve as a quick indicator of problematic graphs within a species, PG-SCUnK offers a more detailed and standardizable alternative. It delivers both a score and an analytical framework, even when evaluating a single graph.

## 4. Conclusion

PG-SCUnK assesses pangenome graph quality through quantifying the representation of single-copy and universal k-mers. This method compares and benchmarks pangenome graphs independently from the source assemblies, thereby enabling to evaluate how well a graph captures the underlying assemblies. PG-SCUnK introduces intuitive metrics that complement traditional graph-level summaries by focusing on sequence representativeness rather than structural properties alone, thereby enhancing the interpretability and utility of graphs for downstream analyses.

### Availability and requirements

Project name: PG-SCUnK

Project home page: https://github.com/cumtr/PG-SCUnK Operating system(s): Linux, MacOS

Programming language: Shell, R Other requirements: KMC v3 License: GNU GPL 3.

Any restrictions to use by non-academics: see License

## Ethics approval and consent to participate

Not applicable

## Consent for publication

Not applicable

## Availability of data and materials

Supplementary data are available online.

The PG-SCUnK workflow is freely available at https://github.com/cumtr/PG-SCUnK. Code used in this article is available at https://github.com/cumtr/PG-SCUnK_paper. The bovine graph produced in this study is available on Zenodo at https://zenodo.org/records/15097551.

Other pangenome graphs used in this study are available at: https://s3-us-west-2.amazonaws-.com/human-pangenomics/index.html?prefix=pangenomes/freeze/freeze1/pggb/chroms/ and https:-//s3-us-west-2.amazonaws.com/human-pangenomics/index.html?prefix=pangenomes/freeze/-freeze1/minigraph-cactus/hprc-v1.1-mc-grch38/hprc-v1.1-mc-grch38.chroms/ for the human pan-genome graphs, https://datadryad.org/stash/dataset/doi:10.5061/dryad.hhmgqnkqb for the finch pangenome graphs, and https://zenodo.org/records/10851548 for the grapevine pangenome graphs.

## Authors’ contributions

TC conceptualized and designed the study with input from HP. TC implemented the method with assistance from ASL. TC conducted the analyses with support from SM and ASL. TC drafted the manuscript, with contributions and revisions from all authors.

## Acknowledgements

Not applicable

## References

Abel HJ, Larson DE, Regier AA et al. Mapping and characterization of structural variation in 17,795 human genomes. Nature 2020;583:83–9.

Auton A, Abecasis GR, Altshuler DM et al. A global reference for human genetic variation. Nature 2015;526:68–74.

Fang B, Edwards SV. Fitness consequences of structural variation inferred from a House Finch pangenome. Proc Natl Acad Sci U S A 2024;121:e2409943121.

Garrison E, Guarracino A, Heumos S et al. Building pangenome graphs. 2023, DOI: 10.1101/2023.04.05.535718.

Garrison E, Sirén J, Novak AM et al. Variation graph toolkit improves read mapping by representing genetic variation in the reference. Nat Biotechnol 2018;36:875–9.

Guarracino A, Heumos S, Nahnsen S et al. ODGI: understanding pangenome graphs. Bioinformatics 2022;38:3319–26.

Günther T, Nettelblad C. The presence and impact of reference bias on population genomic studies of prehistoric human populations. PLOS Genetics 2019;15:e1008302.

Hickey G, Heller D, Monlong J et al. Genotyping structural variants in pangenome graphs using the vg toolkit. Genome Biology 2020;21:35.

Hickey G, Monlong J, Ebler J et al. Pangenome graph construction from genome alignments with Minigraph-Cactus. Nat Biotechnol 2024;42:663–73.

Kokot M, Długosz M, Deorowicz S. KMC 3: counting and manipulating k-mer statistics. Bioinformatics 2017;33:2759–61.

Leonard AS, Crysnanto D, Mapel XM et al. Graph construction method impacts variation representation and analyses in a bovine super-pangenome. Genome Biology 2023;24:124.

Li H, Durbin R. Genome assembly in the telomere-to-telomere era. Nat Rev Genet 2024;25:658–70.

Liao W-W, Asri M, Ebler J et al. A draft human pangenome reference. Nature 2023;617:312–24.

Lin M-J, Iyer S, Chen N-C et al. Measuring, visualizing, and diagnosing reference bias with biastools. Genome Biology 2024;25:101.

Liu Z, Wang N, Su Y et al. Grapevine pangenome facilitates trait genetics and genomic breeding. Nat Genet 2024;56:2804–14.

Martiniano R, Garrison E, Jones ER et al. Removing reference bias and improving indel calling in ancient DNA data analysis by mapping to a sequence variation graph. Genome Biology 2020;21:250.

Milia S, Leonard AS, Mapel XM et al. Taurine pangenome uncovers a segmental duplication upstream of KIT associated with depigmentation in white-headed cattle. Genome Res. 2025: 14;35(4):1041–1052.

Parmigiani L, Garrison E, Stoye J et al. Panacus: fast and exact pangenome growth and core size estimation. Bioinformatics 2024;40:btae720.

Paten B, Eizenga JM, Rosen YM et al. Superbubbles, Ultrabubbles, and Cacti. J Comput Biol 2018;25:649–63.

Shultz AJ, Baker AJ, Hill GE et al. SNPs across time and space: population genomic signatures of founder events and epizootics in the House Finch (Haemorhous mexicanus). Ecol Evol 2016;6:7475–89.

Simão FA, Waterhouse RM, Ioannidis P et al. BUSCO: assessing genome assembly and annotation completeness with single-copy orthologs. Bioinformatics 2015;31:3210–2.

Sirén J, Monlong J, Chang X et al. Pangenomics enables genotyping of known structural variants in 5202 diverse genomes. Science 2021;374:abg8871.

