## supplementary material for "PG-SCUnK: measuring pangenome graph representativeness using single-copy and universal K-mers"

Supplementary information

Supplementary: Material and Methods

### *PG-SCUnK* Implementation

*PG-SCUnK* has been implemented in bash language. It invokes *KMC* (https://github.com/refresh-bio/KMC, Kokot, Długosz and Deorowicz 2017) to efficiently identify single-copy and universal k-mers in assemblies and to classify them as unique, duplicated or split in the pangenome graph.

*PG-SCUnK* operates in three distinct phases.

- *Phase 1: Identification of Single-Copy and Universal k-mers*

In this initial phase, single-copy k-mers are iteratively identified in the input assemblies. First, single-copy k-mers are extracted from a pair of assemblies, and their intersection is computed. This intersected set is then intersected with the single-copy k-mers from a third assembly, and the process is repeated for all assemblies. This iterative approach efficiently identifies single-copy and universal k-mers present in all assemblies while keeping memory requirements low.

- *Phase 2: Extraction and Counting of Graph k-mers*

In the second phase, sequences from the graph nodes are extracted and segmented into k-mers. The frequency of each k-mer is then recorded, providing a detailed profile of the graph's k-mer composition.

- *Phase 3: Comparison and Classification of the k-mers*

In the final phase, the set of universal single-copy k-mers from the input assemblies is compared with the k-mers derived from the graph.

Based on this comparison, k-mers are classified into three categories:

- Unique: k-mers present exactly once in the graph nodes.
- Duplicated: k-mers present more than once in the graph nodes.
- Split: k-mers absent from the graph nodes.

*PG-SCUnK* can handle graphs in the GFAv1 and GFAv2 formats.

### Datasets

Cattle assemblies that were previously published by Milia *et al.* 2024 were provided by the authors. Data can be downloaded from https://www.ebi.ac.uk/ena/browser/view/PRJEB42335.

Human pangenome graphs built by Liao *et al.* 2023 were downloaded from https://s3-us-west-2.amazonaws.com/human-pangenomics/index.html?prefix=pangenomes/freeze/freeze1/pggb/chroms/ for the graph built with *PGGB* (Garrison *et al.* 2024), and from https://s3-us-west-2.amazonaws.com/human-pangenomics/index.html?prefix=pangenomes/freeze/freeze1/minigraph-cactus/hprc-v1.1-mc-grch38/hprc-v1.1-mc-grch38.chroms/ for the graph built with *minigraph cactus* (Hickey *et al.* 2024).

Finch pangenome graphs built by Fang and Edwards 2024 were downloaded from https://datadryad.org/stash/dataset/doi:10.5061/dryad.hhmgqnkqb.
All autosomal chromosomes were considered for our analyses except chromosome 16 for which the naming scheme of the graph indicated a distinct *PGGB* run and chromosomes 31 to 39 which were substantially smaller (less than 5Mbp) than the other chromosomes.

Grapevine pangenome graphs built by Liu *et al.* 2024 were downloaded from https://zenodo.org/records/10851548.

For each organism, reference genome and genome statistics (Number of chromosomes, total size, size of each chromosome, GC content) were retrieved from the NCBI:
Cattle: https://www.ncbi.nlm.nih.gov/datasets/genome/GCF_002263795.3/
Human: https://www.ncbi.nlm.nih.gov/datasets/genome/GCF_000001405.40/
Finch: https://www.ncbi.nlm.nih.gov/datasets/genome/GCF_027477595.1/
Grapevine: https://www.ncbi.nlm.nih.gov/datasets/genome/GCF_030704535.1/

#### Cattle pangenome graph building

All cattle graphs were built with *PGGB* (Garrison *et al.* 2024). The *PGGB* workflow (v0.6.0) was executed using *wfmash* (v0.21.0-318-gb160f53) (Marco-Sola *et al.* 2021), *seqwish* (v0.7.10-0-g75e807c) (Garrison and Guarracino 2023), *smoothxg* (v0.8.0-2-ge93c623) (https://github.com/pangenome/-smoothxg), *gfaffix* (v0.1.5) (https://github.com/marschall-lab/GFAffix, Liao *et al.* 2023), and *odgi* (v0.8.6-0-ge647844f) (Guarracino *et al.* 2022).

The default cattle graphs for the autosomes were built with a percent-identity score (-p) of 98%, a segment length (-s) of 75k and a min-match-length (-k) of 31.

The same *PGGB* workflow was used to investigate the impact of the p, k, and s parameters on the *PG-SCUnK* scores. All analyses were performed for bovine chromosome 13. For each parameter varying, the two other parameters were set to the values used for the default graphs. We built 6 graphs with different values for the segment-length (-s): 1k, 5k, 10k, 25k, 50k and 100k (with-k 31 and-p 98). We built 12 graphs with different values of the percent-identity score (-p): 70, 75, 80, 85, 90, 95, 97, 97.5, 98, 98.5, 99, 99.5 (with-s 75k and-k 31). We built 9 graphs with different values for the min-match-length (-k) parameter: 11, 21, 31, 41, 51, 111, 211, 311, 411 (with-s 75k and-p 98).

#### Graph statistics

Total length and number of nodes and edges were extracted from the graphs using *odgi* (v0.8.6-0-ge647844f) (Guarracino *et al.* 2022). First, *gfa* graphs were transformed into *og* files using the *odgi build* command (-P -O -s parameters). Graph statistics were then extracted with the *odgi stats* command (-S-ps parameters).

#### Running PG-SCUnK

*PG-SCUnK* was run on all the graphs in two steps. First, the companion workflow *GFA2HaploFasta.bash* was run to extract all the assemblies from a given graph through calling the *paths* command from *odgi* (v0.8.6-0-ge647844f). Then, *PG-SCUnK* was run on those assemblies with a k-mer size (-k) of 100.

#### Impact of PG-SCUnK k-mer size

To measure the impact of k-mer size on the proportion of the k-mers considered as SCUnKs as well as the *PG-SCUnK* scores, we ran *PG-SCUnK* for different values of the –k parameter including 31, 51, 71 ,100 [default], 111 and 211.
We estimated the total number of unique k-mers in all the chromosomes of the different species using the reference genome of each species using *KMC* v3.2.4 (Kokot, Długosz and Deorowicz 2017).

#### Short reads simulation and alignment

Short reads (150 bp) were simulated from each assembly using *wgsim* (v 0.3.1-r13, https://github.com/lh3/wgsim). The number of reads simulated per assembly was determined to produce a coverage of 15X for each assembly. *wgsim* was run with the parameters-1 150-2 150-e 0-r 0-R 0-X 0, and only forward reads were retrieved for mapping.

We built an index for the chromosome level graphs for all species using *vg* (v1.60.0 "Annicco") (Sirén *et al.* 2021). We used the *vg autoindex* command with the --*workflow giraffe* parameter. Indexing failed for some chromosomes due to exceeding memory or runtime (5 threads, 250 GB RAM, 300 hours). Affected chromosomes were excluded from downstream analyses. These included chromosomes 1, 4, and 7 in the finch pangenome, chromosomes 2, 12, and 14 in grapevine, and chromosomes 1 and 13 in humans. For successfully indexed chromosomes, short-read mapping was performed using *vg giraffe* (v1.60.0 "Annicco") with default parameters. Mapping failed for some chromosomes due to exceeding runtime (50 threads, 100 hours). Affected chromosomes were excluded from downstream analyses. Mapping statistics were extracted from the GAM files using a custom script (*GetStats.py*), available at https://github.com/cumtr/PG-SCUnK_paper.

Supplementary: Figures and Tables

Table S1 – Summary statistics of the pan-genome graphs used in this study.

| ORGANISM | TOTAL SIZE (BP) | # OF NODES | # OF EDGES | MEAN NODE SIZE (BP) | MEAN EDGES / NODE |
|---|---|---|---|---|---|
| **CATTLE** | 5’136’671’978 | 168’192’544 | 231’479’631 | 30.54 | 1.3763 |
| **HUMAN (PGGB)** | 8’120’641’583 | 107’104’689 | 149’426’113 | 75.82 | 1.3951 |
| **HUMAN (MC)** | 12’321’448’492 | 77’754’446 | 107’832’366 | 158.47 | 1.3868 |
| **FINCH** | 2’723’680’256 | 196’241’920 | 276’077’085 | 13.88 | 1.4068 |
| **GRAPEVINE** | 1’525’722’021 | 109’877’636 | 155’276’528 | 13.89 | 1.4132 |

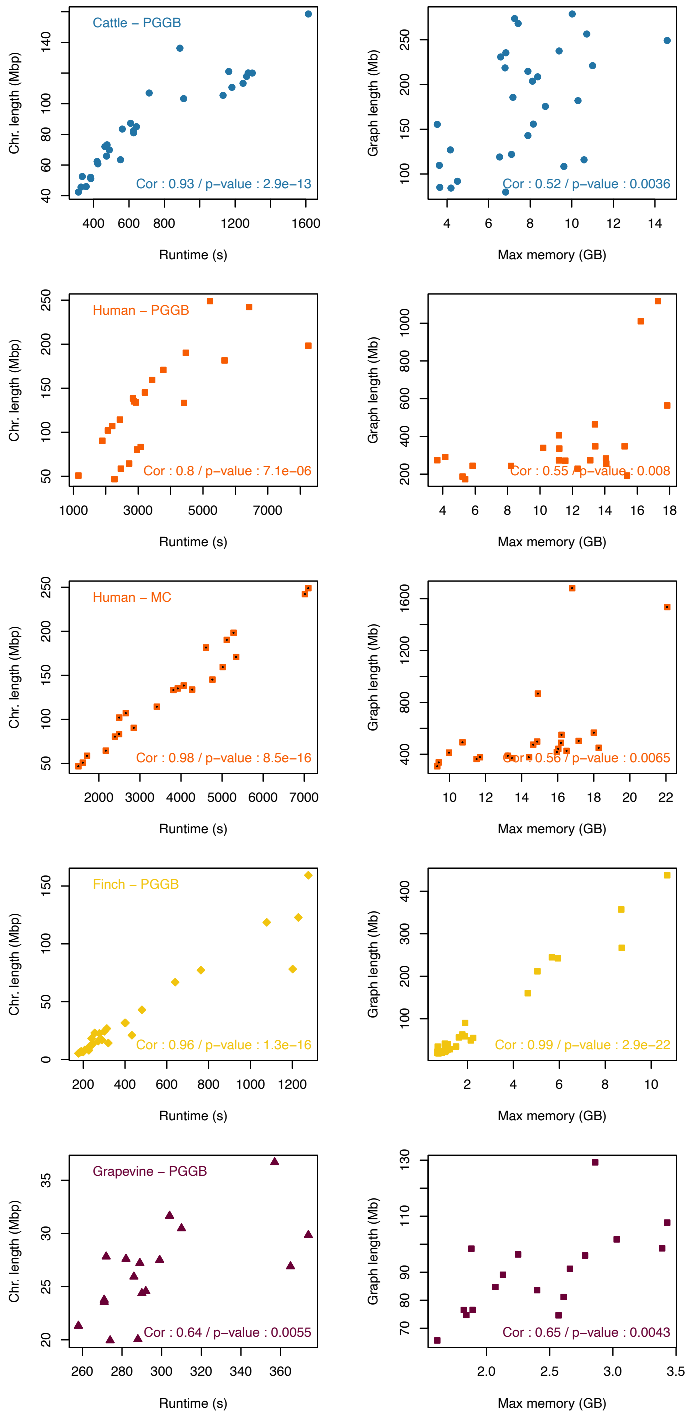

**Figure S1 – Runtime and memory usage of PG-SCUnK correlated with chromosome length and graph size respectively**. Left panels: Scatterplots between runtime and chromosomal size for all the pangenome graphs. Right panels: Scatterplots between the maximum memory usage and the graph size for the different pangenome graphs. The top left legend in plots specify which pangenome is represented in the row. The bottom right legend presents the Pearson's correlation coefficient between the two variables and the associated p-value.

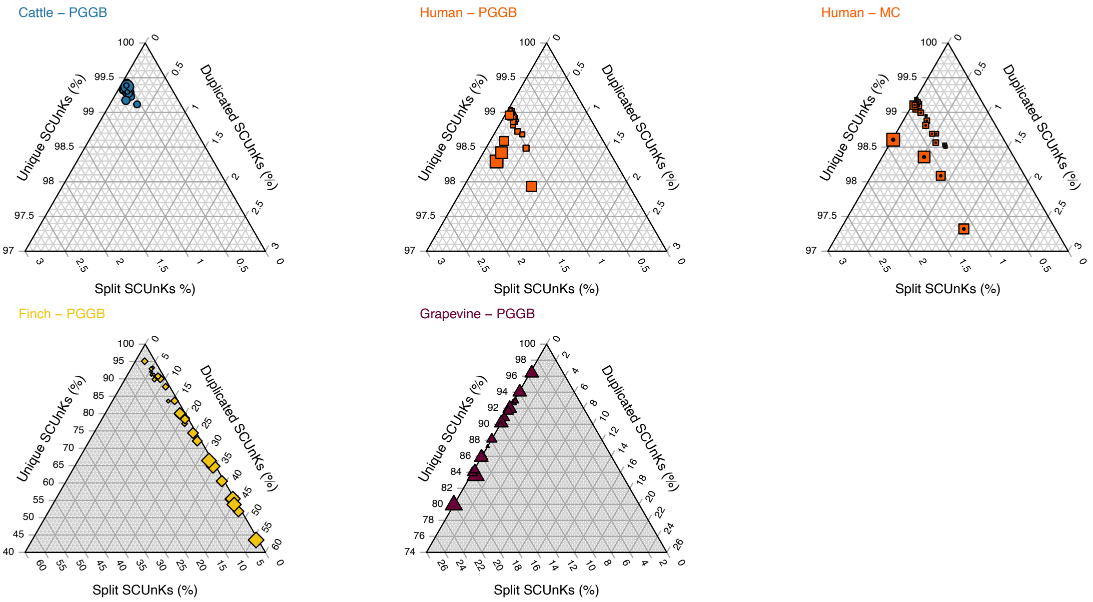

**Figure S2 – Ternary plots showing the relation between the PG-SCUnK across different organisms.** PG-SCUnK scores for all chromosome-level pangenome graphs. Dot size depicts GC content of the chromosome, with larger dots indicating higher GC content.

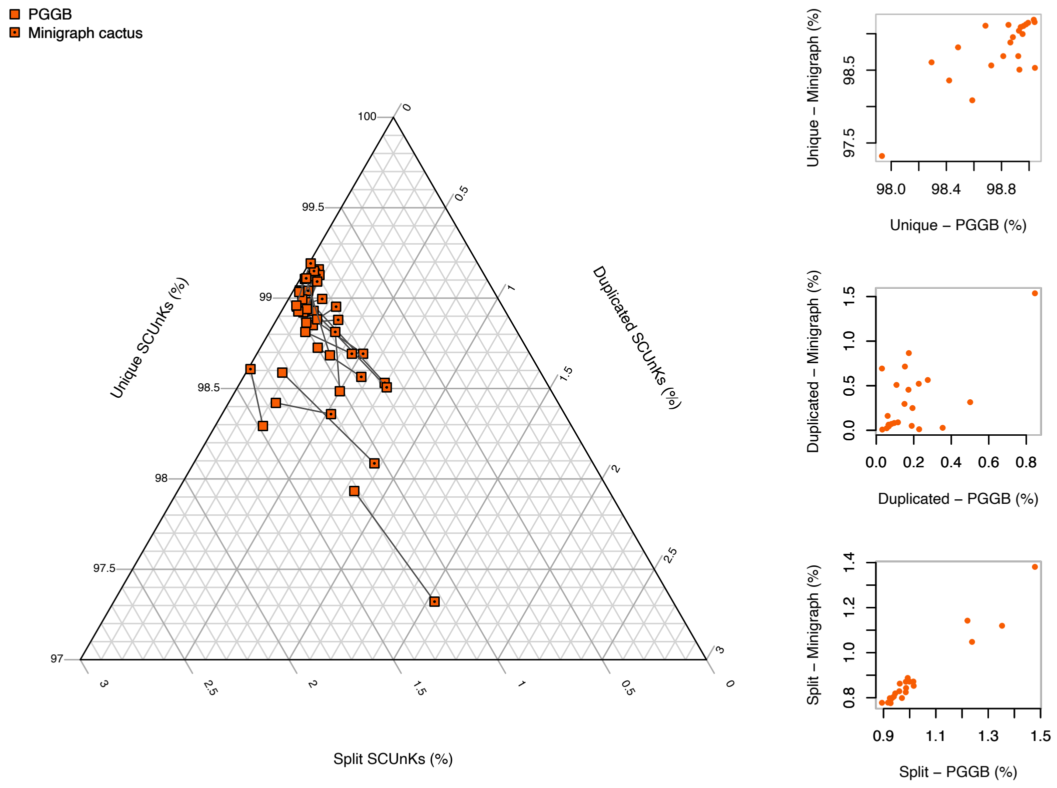

**Figure S3 – PG-SCUnK scores for human pangenome graphs built with two different software**. Full squares represent chromosome-level pangenome graphs built with PGGB, squares with a dot inside represent the chromosome-level pangenome graphs built with Mingraph cactus. Symbols linked by segments represent the same chromosome. Right panels: Scatterplots between the percentage of unique (top), duplicated (center) and split k-mers (bottom) from the two software.

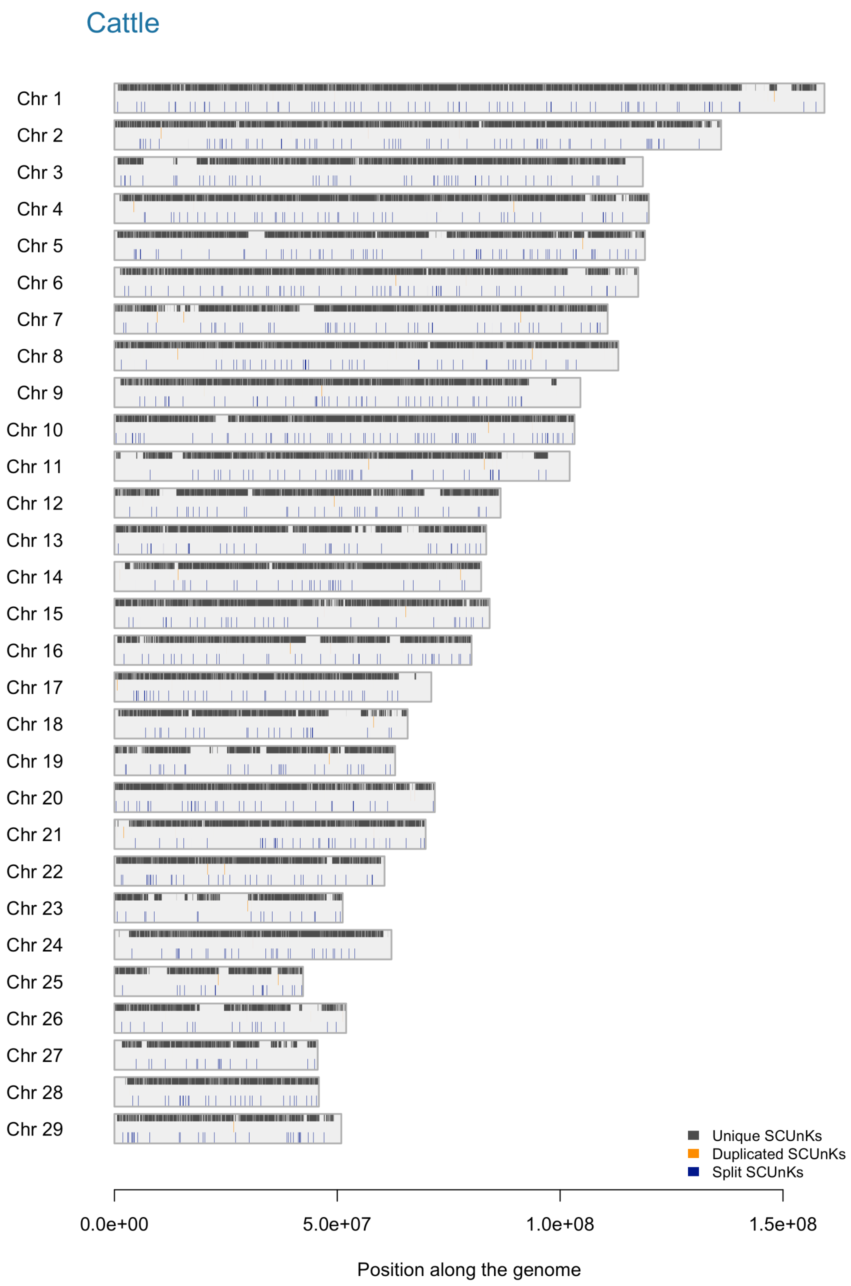

**Figure S4 – Distribution of the unique, duplicated and split SCUnKs along an examined cattle genome (ARS_UCD1.2)**. Each rectangle represents a chromosome. Within each chromosome, colored ticks indicate the locations of unique, duplicated, and split SCUnKs.

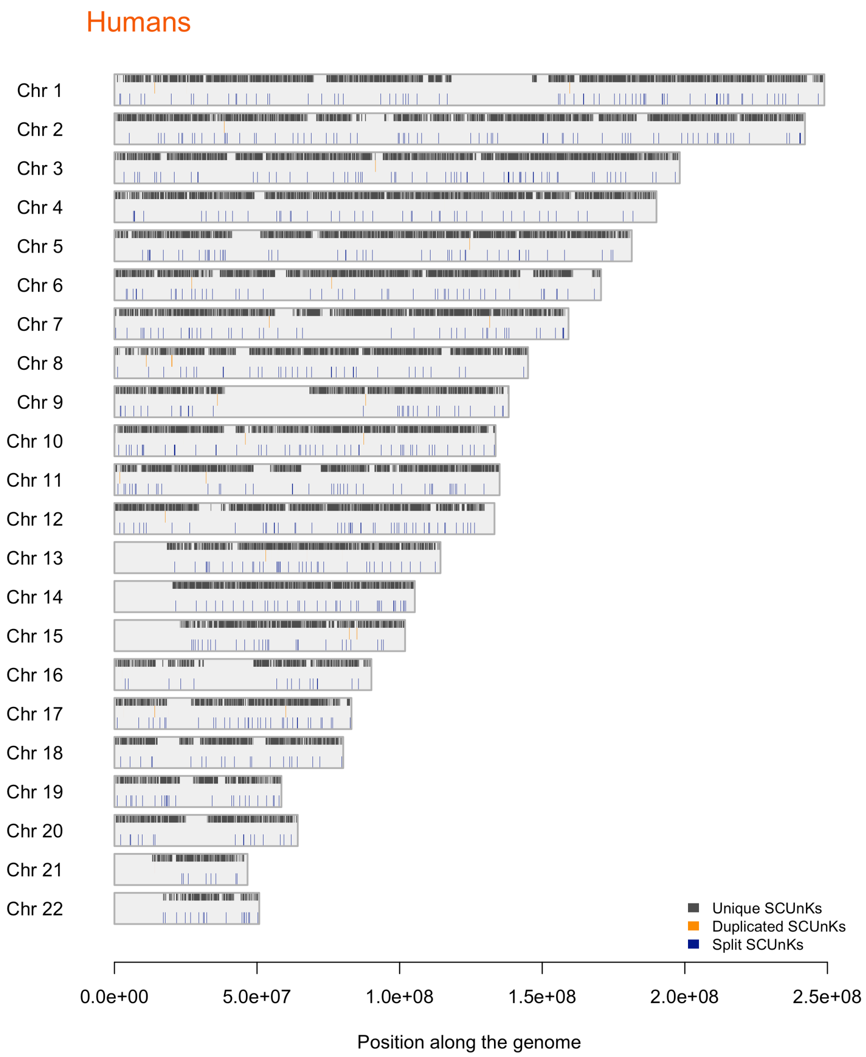

**Figure S5 – Distribution of the unique, duplicated and split SCUnKs along an examined human genome (GRCh38)**. Each rectangle represents a chromosome. Within each chromosome, colored ticks indicate the locations of unique, duplicated, and split SCUnKs.

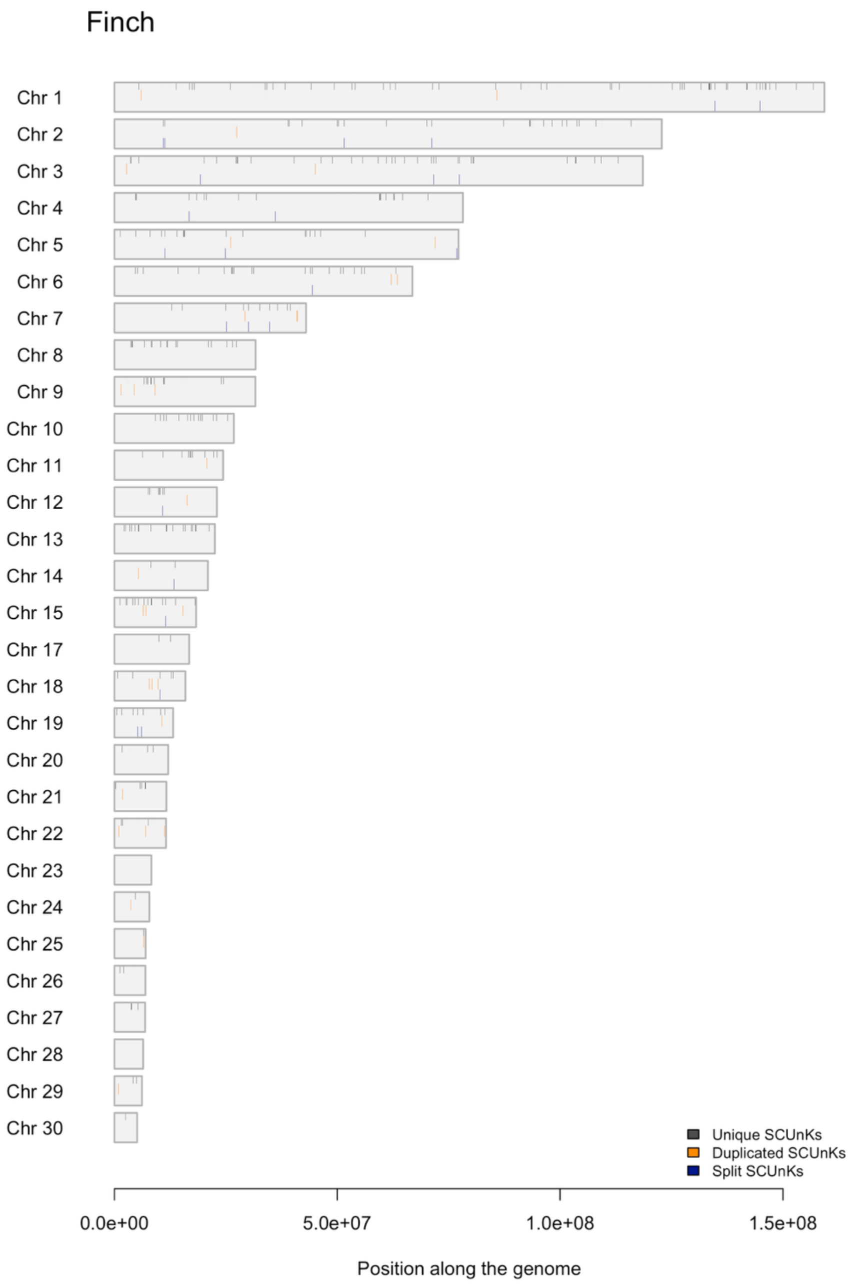

**Figure S6 – Distribution of the unique, duplicated and split SCUnKs along an examined finch genome (bHaeMex1).** Each rectangle represents a chromosome. Within each chromosome, colored ticks indicate the locations of unique, duplicated, and split SCUnKs.

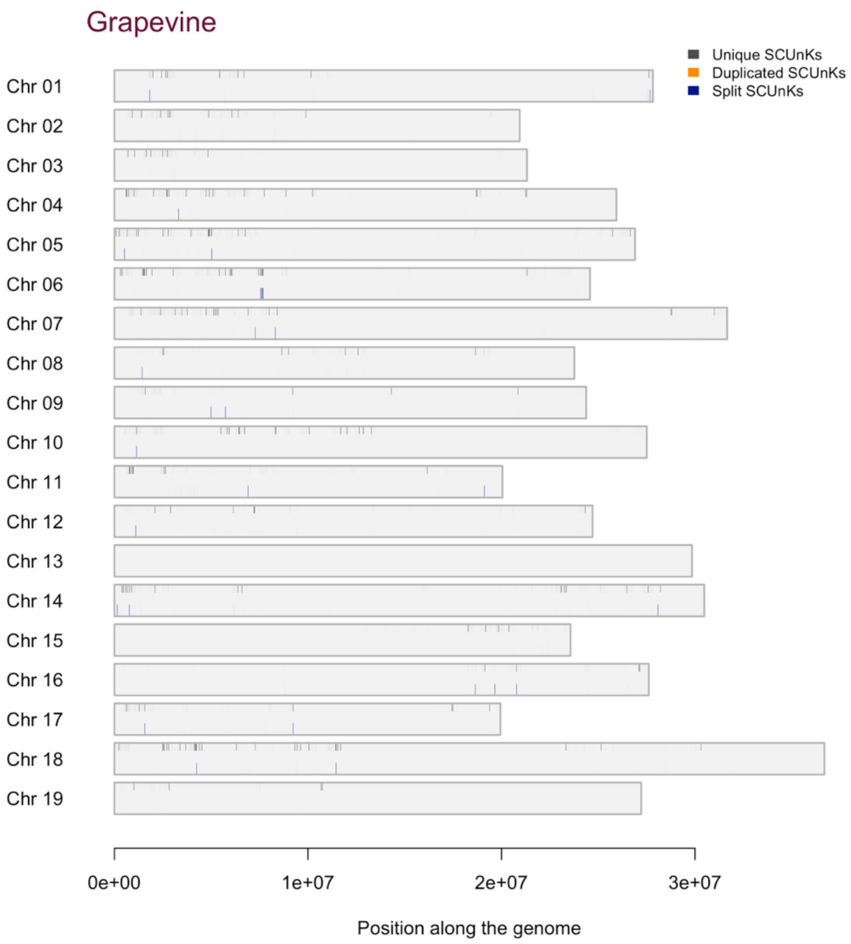

**Figure S7 – Distribution of the unique, duplicated and split SCUnKs along an examined grapevine genome (PN)**. Each rectangle represents a chromosome. Within each chromosome, colored ticks indicate the locations of unique, duplicated, and split SCUnKs.

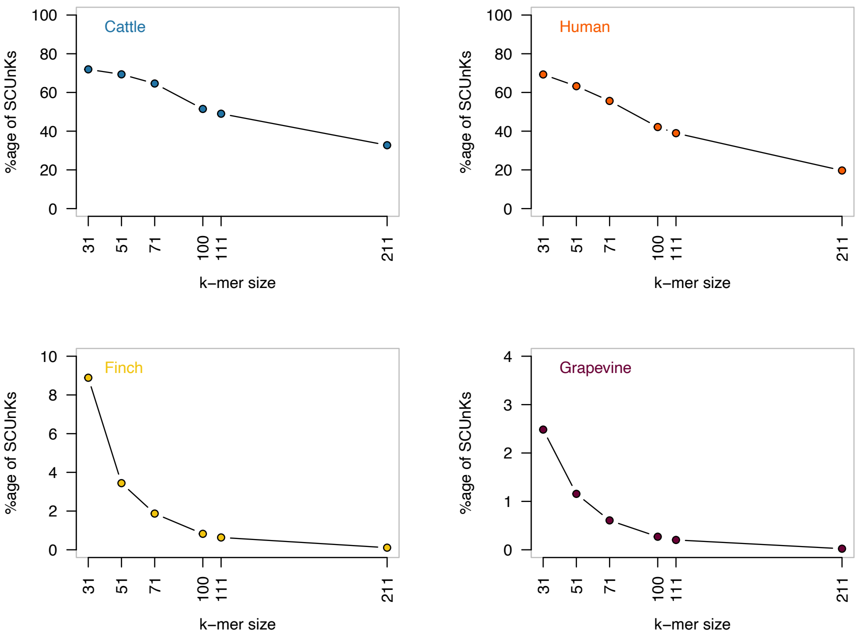

**Figure S8 – Percentage of the total k-mers that are SCUnKs in the reference genome for different k-mer size (-k parameter)**. Each panel depicts a different organism.

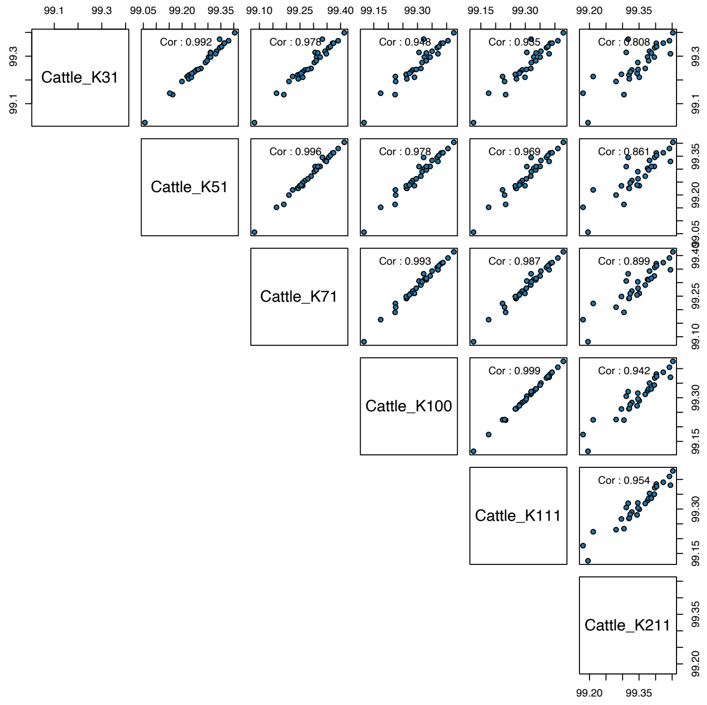

**Figure S9 – Correlation between the PG-SCUnK scores for different values of k in cattle**. Each dot represents the proportion of unique SCUnKs for a given chromosome.

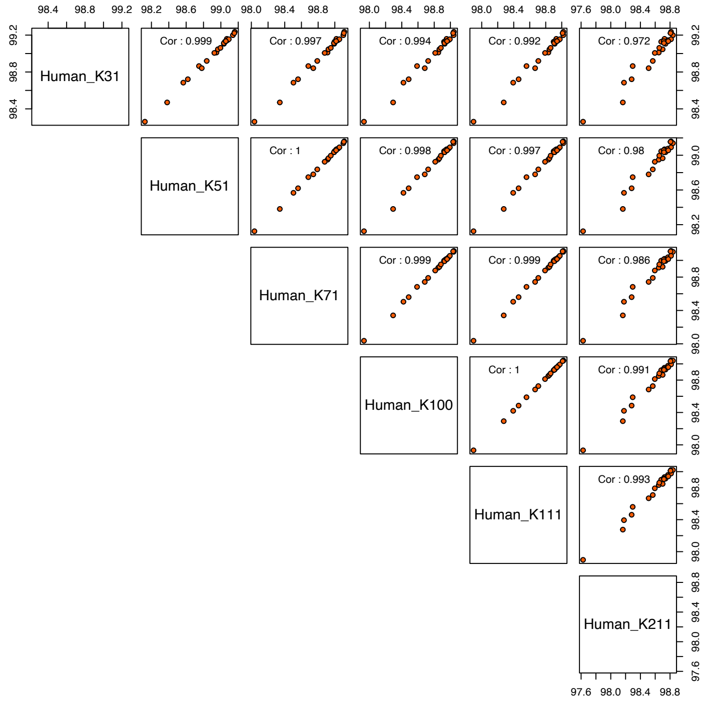

**Figure S10 – Correlation between the PG-SCUnK scores for different values of k in human**. Each dot represents the proportion of unique SCUnKs for a given chromosome. The results presented here are for the human pangenome graph build with PGGB.

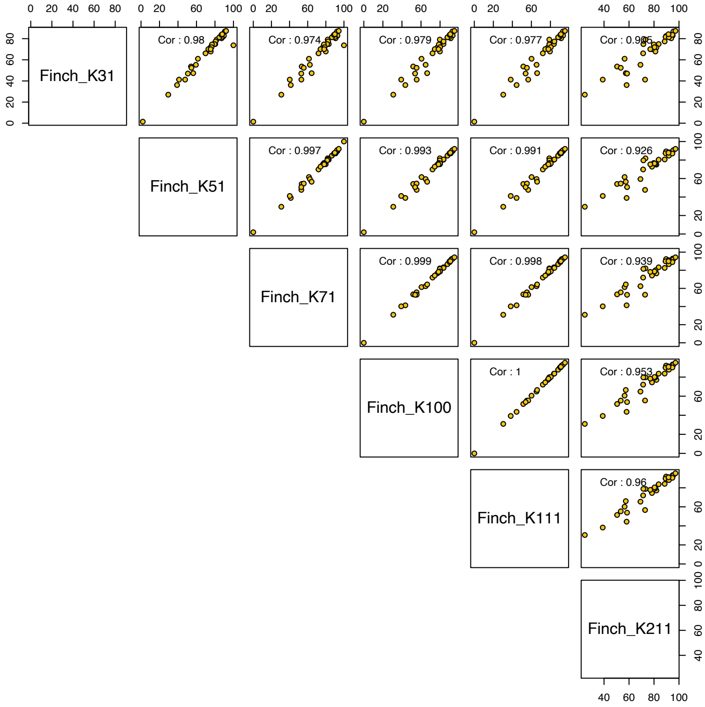

**Figure S11 – Correlation between the PG-SCUnK scores for different values of k in finch**. Each dot represents the proportion of unique SCUnKs for a given chromosome.

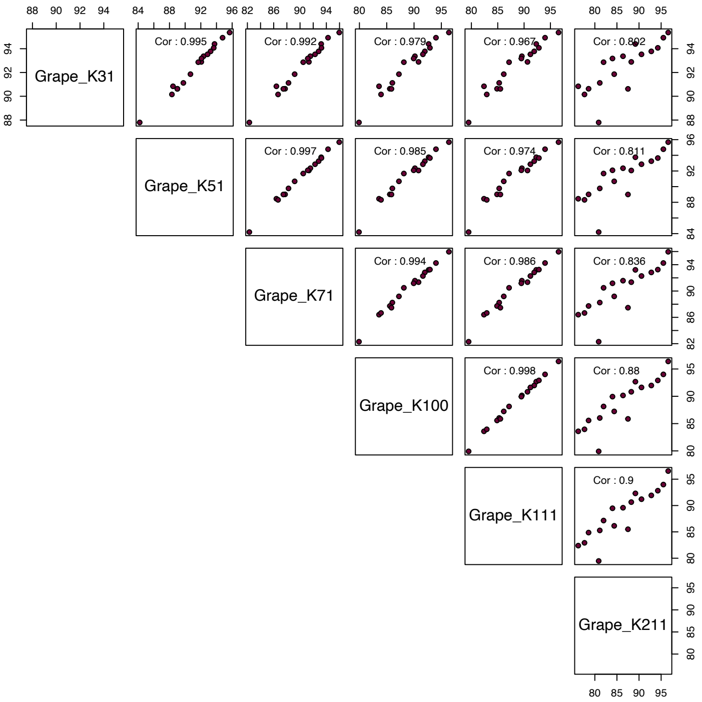

**Figure S12 – Correlation between the PG-SCUnK scores for different values of k in grapevine**. Each dot represents the proportion of unique SCUnKs for a given chromosome.

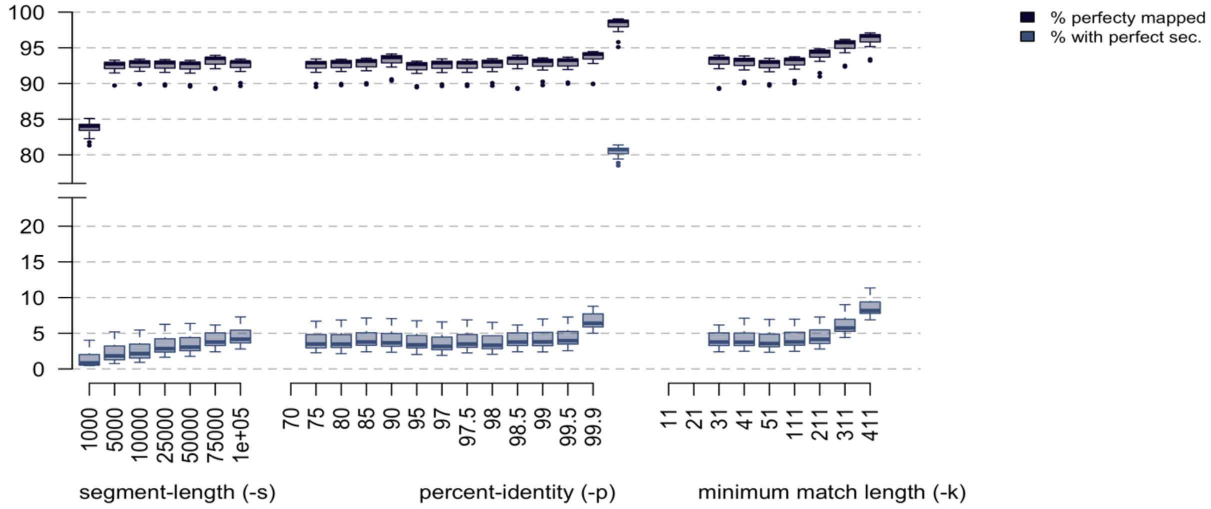

**Figure S13 – Mapping quality of simulated reads against the Cattle pangenome graphs build with different parameter**. Each boxplot depict the distribution of the proportion of reads mapped with a mapping quality of 60 (darker color), with the proportion of them having at least one secondary mapping as good as the first (i.e. both have a mapping score of 60).

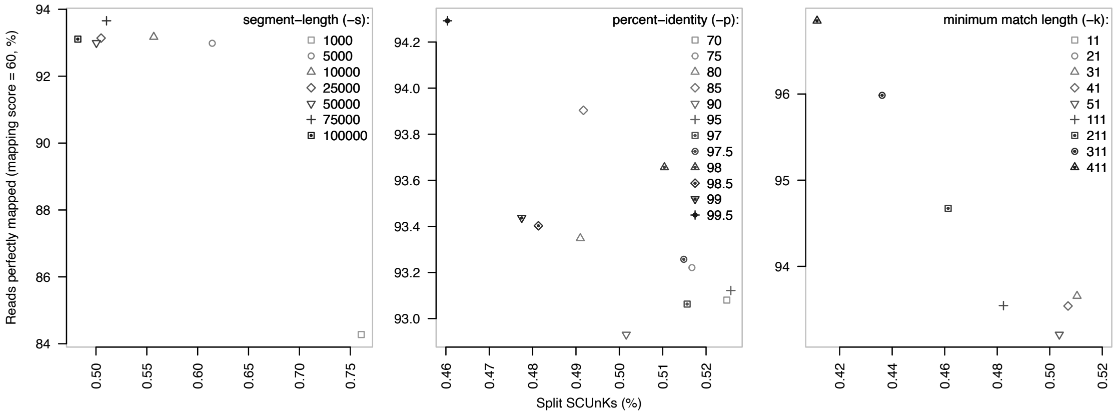

**Figure S14 – Negative relation between the proportion of mapped read (mapping quality = 60) and the rate of split SCUnKs**. Each panel depicts a different parameter range used during the building of a cattle pangenome graph for chromosome 13. For each varying parameter, the two others were fixed to the values used for the entire genome (fixed parameters:-s 75k,-p 98, k 31).

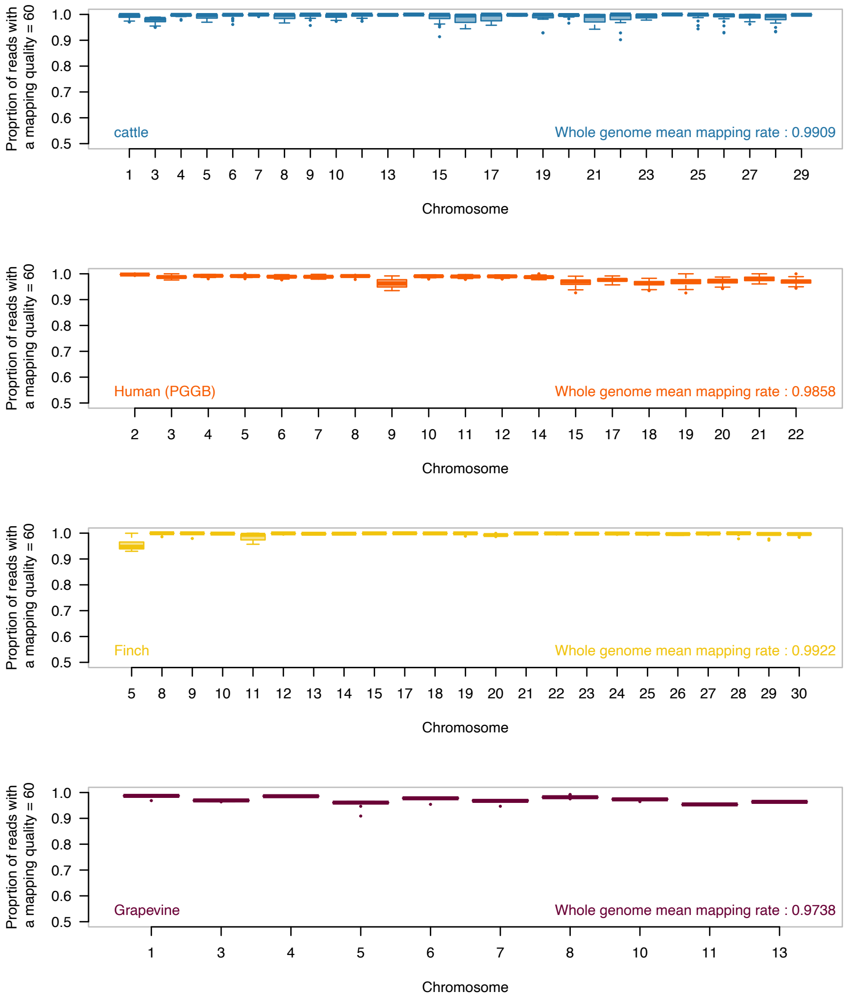

**Figure S15 – Mapping quality of simulated reads against pangenome graphs.** Proportion of reads with a primary mapping quality of 60. Each panel represents the result for one organism, with one boxplot per chromosome. Missing chromosomes failed either indexing or mapping, see material and method section above for details.

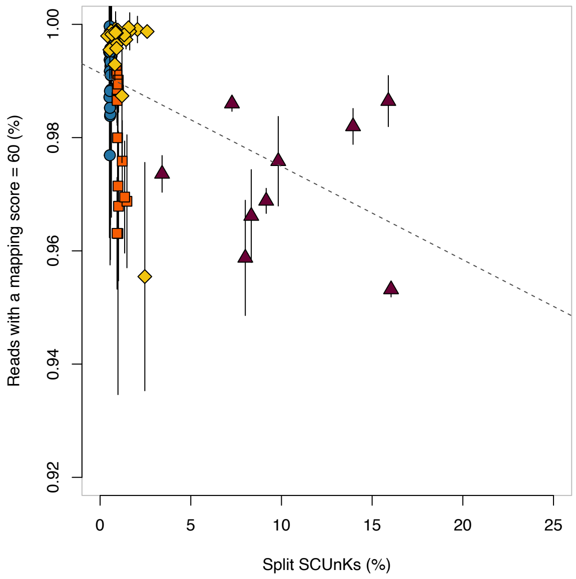

**Figure S16 – Relation between the PG-SCUnK split score and the proportion of reads with a perfect alignment (mapping quality = 60) across the different organisms.** Each dot represents the mean mapping rate for a chromosome (circle for Cattle, square for Human, diamond for the Finch and triangle for the Grapevine) with the bar presenting the standard deviation around this mean.

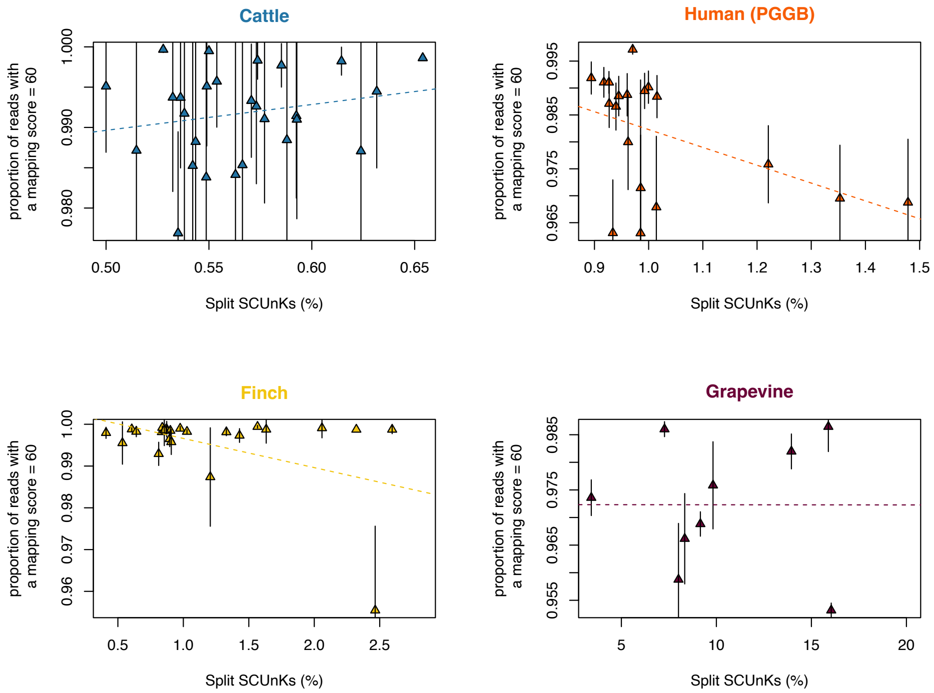

**Figure S17 – Relation between the PG-SCUnK split score and the proportion of reads with a perfect alignment (mapping quality = 60) within the different organisms.** Top left plot is for the cattle graphs, top right for the human graphs, bottom left is for the finch graphs, and bottom right is for the grapevine graphs. For each organism, each dot represents the mean mapping rate for a chromosome with the bar presenting the standard deviation around this mean.

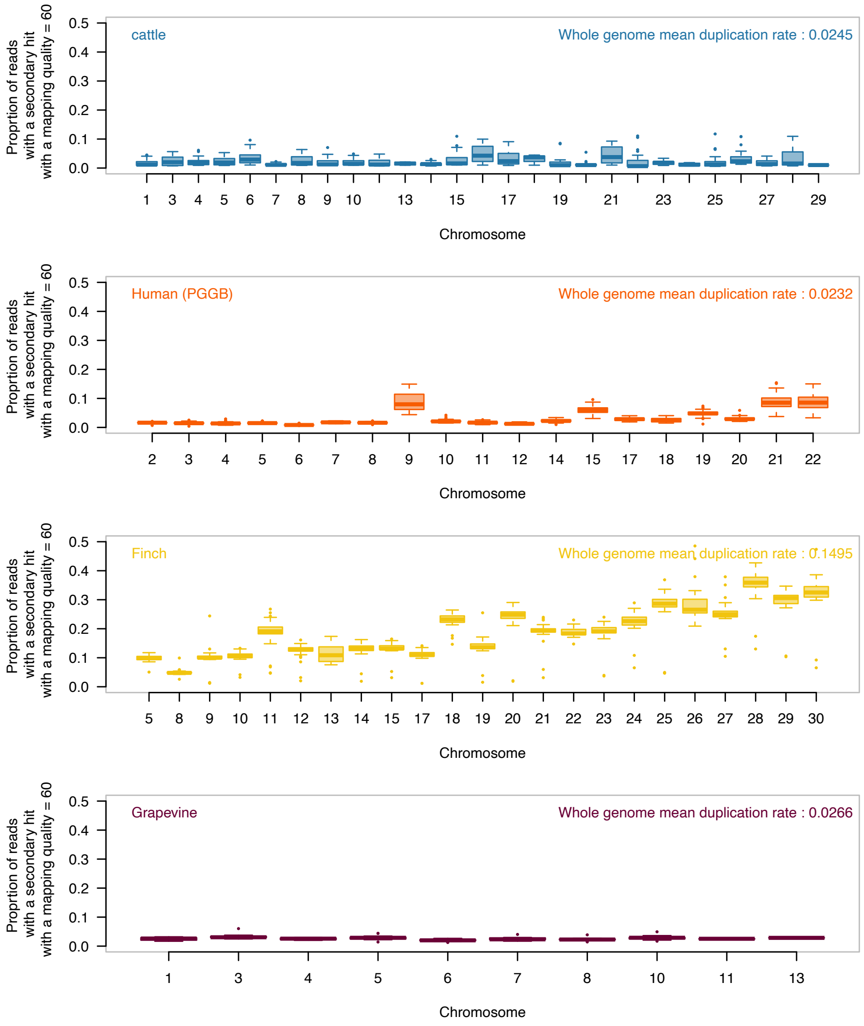

**Figure S18 – Proportion of simulated reads with a perfect secondary alignment (mapping quality = 60).** Each panel presents the result for one organism, with one boxplot per chromosomes. Missing chromosomes failed either indexing or mapping, see material and method section above for details.

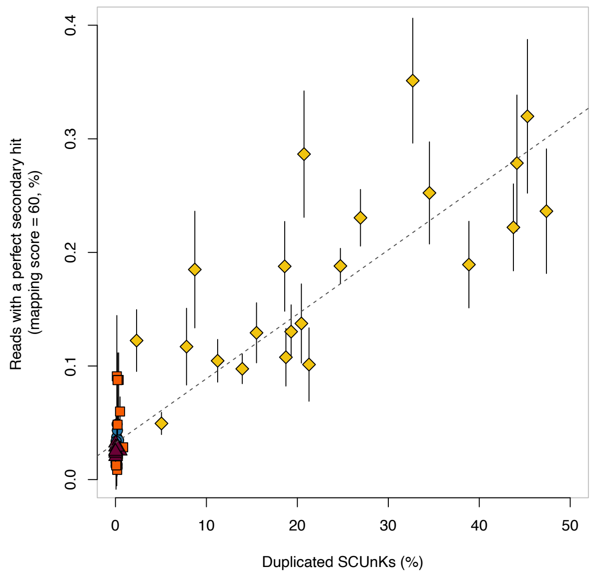

**Figure S19 – Relation between the PG-SCUnK split score and the proportion of reads with a perfect alignment (mapping quality = 60) across the different organisms.** Each dot represents the mean mapping rate for a chromosome (circle for Cattle, square for Human, diamond for the Finch and triangle for the Grapevine) with the bar presenting the standard deviation around this mean.

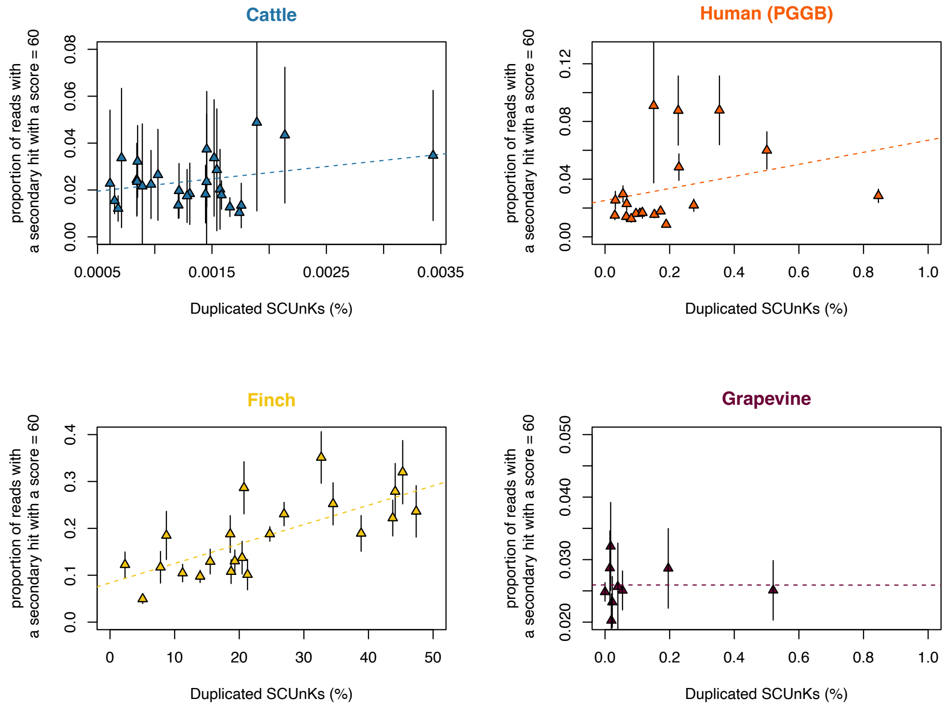

**Figure S20 – Relation between the PG-SCUnK duplication score and the proportion of reads with a perfect secondary alignment (mapping quality = 60) within the different organisms.** Top left plot is for the cattle graphs, top right for the human graphs, bottom left is for the finch graphs, and bottom right is for the grapevine graphs. For each organism, each dot represents the mean mapping rate for a chromosome with the bar presenting the standard deviation around this mean.

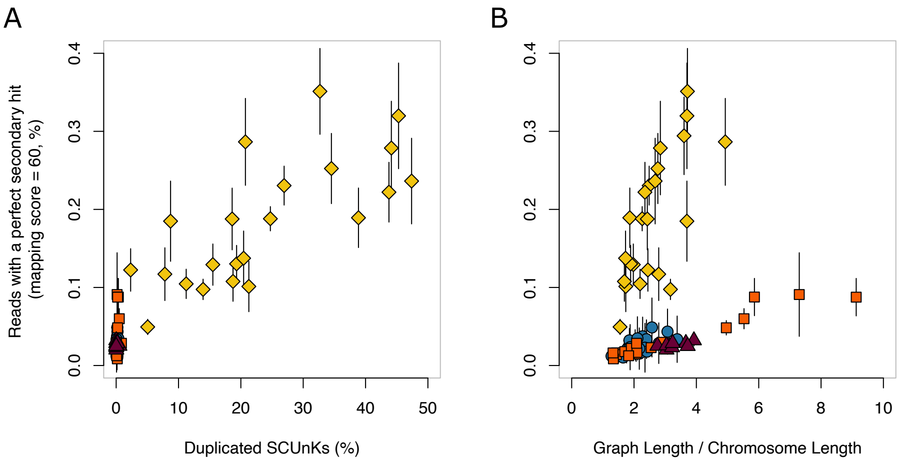

**Figure S21 – Relation between the PG-SCUnK duplication score and the ratio (Graph length) / (assembly length) with the proportion of reads with a perfect secondary alignment (mapping quality = 60) across the different organisms**. (A) Relation between the PG-SCUnK duplication score and the proportion of reads with a perfect secondary alignment. (B) Relation between the ratio (Graph length) / (assembly length) and the proportion of reads with a perfect secondary alignment**.** In the two graphs, Each dot represents the mean mapping rate for a chromosome (circle for Cattle, square for Human, diamond for the Finch and triangle for the Grapevine) with the bar presenting the standard deviation around this mean.

### References

Fang B, Edwards SV. Fitness consequences of structural variation inferred from a House Finch pangenome. *Proc Natl Acad Sci U S A* 2024;**121**:e2409943121.

Garrison E, Guarracino A. Unbiased pangenome graphs. *Bioinformatics* 2023;**39**:btac743.

Garrison E, Guarracino A, Heumos S *et al.* Building pangenome graphs. *Nat Methods* 2024;**21**:2008–12.

Guarracino A, Heumos S, Nahnsen S *et al.* ODGI: understanding pangenome graphs. *Bioinformatics* 2022;**38**:3319–26.

Hickey G, Monlong J, Ebler J *et al.* Pangenome graph construction from genome alignments with Minigraph-Cactus. *Nat Biotechnol* 2024;**42**:663–73.

Kokot M, Długosz M, Deorowicz S. KMC 3: counting and manipulating k-mer statistics. *Bioinformatics* 2017;**33**:2759–61.

Liao W-W, Asri M, Ebler J *et al.* A draft human pangenome reference. *Nature* 2023;**617**:312–24.

Liu Z, Wang N, Su Y *et al.* Grapevine pangenome facilitates trait genetics and genomic breeding. *Nat Genet* 2024;**56**:2804–14.

Marco-Sola S, Moure JC, Moreto M *et al.* Fast gap-affine pairwise alignment using the wavefront algorithm. *Bioinformatics* 2021;**37**:456–63.

Milia S, Leonard A, Mapel XM *et al.* Taurine pangenome uncovers a segmental duplication upstream of KIT associated with depigmentation in white-headed cattle. *Genome Res* 2024:gr.279064.124.

Sirén J, Monlong J, Chang X *et al.* Pangenomics enables genotyping of known structural variants in 5202 diverse genomes. *Science* 2021;**374**:abg8871.
